# MicroRNAs and mRNA Regulatory Network of Parenchymal Hematoma after Endovascular Mechanical Reperfusion for Acute Ischemic Stroke in Rat

**DOI:** 10.1101/2024.02.01.578357

**Authors:** Jin-Kun Zhuang, Zhong-Run Huang, Wang Qin, Chang-Luo Li, Qi Li, Chun Xiang, Yong-Hua Tuo, Zhong Liu, Qian-Yu Chen, Zhong-Song Shi

## Abstract

Hemorrhagic transformation after endovascular thrombectomy predicts poor outcomes in acute ischemic stroke with large vessel occlusion. The roles of microRNAs in the pathogenesis of parenchymal hematoma (PH) are still unclear. This study aims to investigate the microRNA and mRNA regulatory network associated with PH after mechanical reperfusion in the animal stroke model and oxygen-glucose deprivation/reoxygenation (OGD/R) model. Twenty-five microRNAs were assessed in the reperfusion-induced hemorrhage model in rats with hyperglycemic conditions receiving 5-hour middle cerebral artery occlusion. Thirteen down-regulated microRNAs (miRNA-29a-5p, miRNA-29c-3p, miRNA-126a-5p, miRNA-132-3p, miRNA-136-3p, miRNA-142-3p, miRNA-153-5p, miRNA-218a-5p, miRNA-219a-2-3p, miRNA-369-5p, miRNA-376a-5p, miRNA-376b-5p, miRNA-383-5p) and one up-regulated microRNA (miRNA-195-3p) were found in rat peri-infarct with PH. Ten of these 14 PH-related microRNAs were significantly differentially expressed in at least two of five models of neuron, astrocyte, microglia, BMEC, and pericyte after OGD/R, consistent with the animal model results. Thirty-one predicted hub target genes were significantly differentially expressed in rat peri-infarct with PH. Forty-nine microRNA-mRNA regulatory axes of PH were revealed, which were related to the mechanisms of oxidative stress, apoptosis, immune, and inflammation. Simultaneously differentially expressed microRNAs and related genes in several cells of the neurovascular unit may serve as valuable targets for PH after endovascular thrombectomy in acute ischemic stroke.

## Introduction

Endovascular thrombectomy (EVT) alone or intravenous alteplase plus EVT can provide clinical benefit in patients with acute ischemic stroke (AIS) due to large vessel occlusion shown in the randomized controlled trials [1–6]. Half of patients with AIS receiving EVT have favorable outcomes. Hemorrhagic transformation (HT) occurs in one-third of patients with AIS receiving EVT alone or intravenous alteplase plus EVT, even in half of the patients having EVT beyond 6 hours after stroke [7–9]. Symptomatic HT, one of the predictors of poor outcome, was recently reduced, accompanied by the advancement of EVT. However, parenchymal hematoma (PH) is still associated with functional dependence after EVT [10–12].

The blood-brain barrier (BBB) dysfunction is vital in cerebral ischemia-reperfusion injury and HT [13–15]. Early BBB disruption predicts HT and poor outcomes in acute stroke patients receiving EVT [16–18]. Non-coding RNA can contribute to cerebral ischemia-reperfusion injury and act as non-invasive biomarkers of diagnosis and prognosis for HT in patients with acute stroke [19–22]. In previous studies, we showed the preliminary results of the microRNA expression profile in rat ischemic hemispheres with HT after mechanical reperfusion using a microRNA array. Nicotinamide adenine dinucleotide phosphate oxidase inhibitor reduced HT after endovascular mechanical reperfusion and altered the levels of several microRNAs [23, 24]. The purpose of this study is to investigate the microRNA and mRNA regulatory network associated with PH after mechanical reperfusion in vivo animal model and in-vitro oxygen-glucose deprivation/reoxygenation (OGD/R) model of neuron, astrocyte, microglia, brain microvascular endothelial cell (BMEC), and pericyte.

## Materials and Methods

### MCAO model with hemorrhagic transformation after mechanical reperfusion

Adult male Sprague-Dawley rats (250-280g, 8-10 weeks) were used to create the model of reperfusion-induced HT with hyperglycemic conditions. The intraluminal filament technique was performed to have the middle cerebral artery occlusion (MCAO) for 5 hours and then receive reperfusion after 19 hours, as described in our previous study [23, 24]. We prolonged the MCAO time for 5 hours, resulting in severe cerebral ischemia, and removed the filament to allow instant reperfusion. This model has achieved cerebral blood flow and pathophysiological alteration upon reperfusion, similar to mechanical thrombectomy in humans [25]. Anesthesia was induced with 5% isoflurane in a chamber with 70% nitrogen and 30% oxygen and maintained with 2% isoflurane using a mask. All rats received intraperitoneal injection of 50% dextrose (6ml/kg) 15 minutes before MCAO to induce the condition of hyperglycemia with a glucose level larger than 16.7 mmol/L. Subsequently, 50% dextrose (1.5ml/kg) was injected intraperitoneally at 1, 2, 3, and 4 hours after MCAO to maintain the hyperglycemic state.

Fifty-two rats with hyperglycemic states were in the HT group (n=42) and the sham-operated group (n=10). The sham-operated hyperglycemic rats underwent the same surgical procedures without inserting the filament. The rats were sacrificed 19 hours after removing the filament (24 hours after the onset of ischemia). Rat brains were harvested for measurements. HT was classified into hemorrhagic infarction and parenchymal hemorrhage, according to previous studies [23, 24]. The coronal brain slices assessed macroscopic HT. Hemorrhagic infarction was defined as small or more confluent petechiae within the infarcted area or at the borders of the ischemic area. PH was shown as a large area of blood within the infarcted area (Figure 1). The Institutional Animal Research Committee of Sun Yat-sen Memorial Hospital of Sun Yat-sen University approved the experimental protocol in accordance with policies set by the National Institutes of Health guidelines.

**Figure 1.**
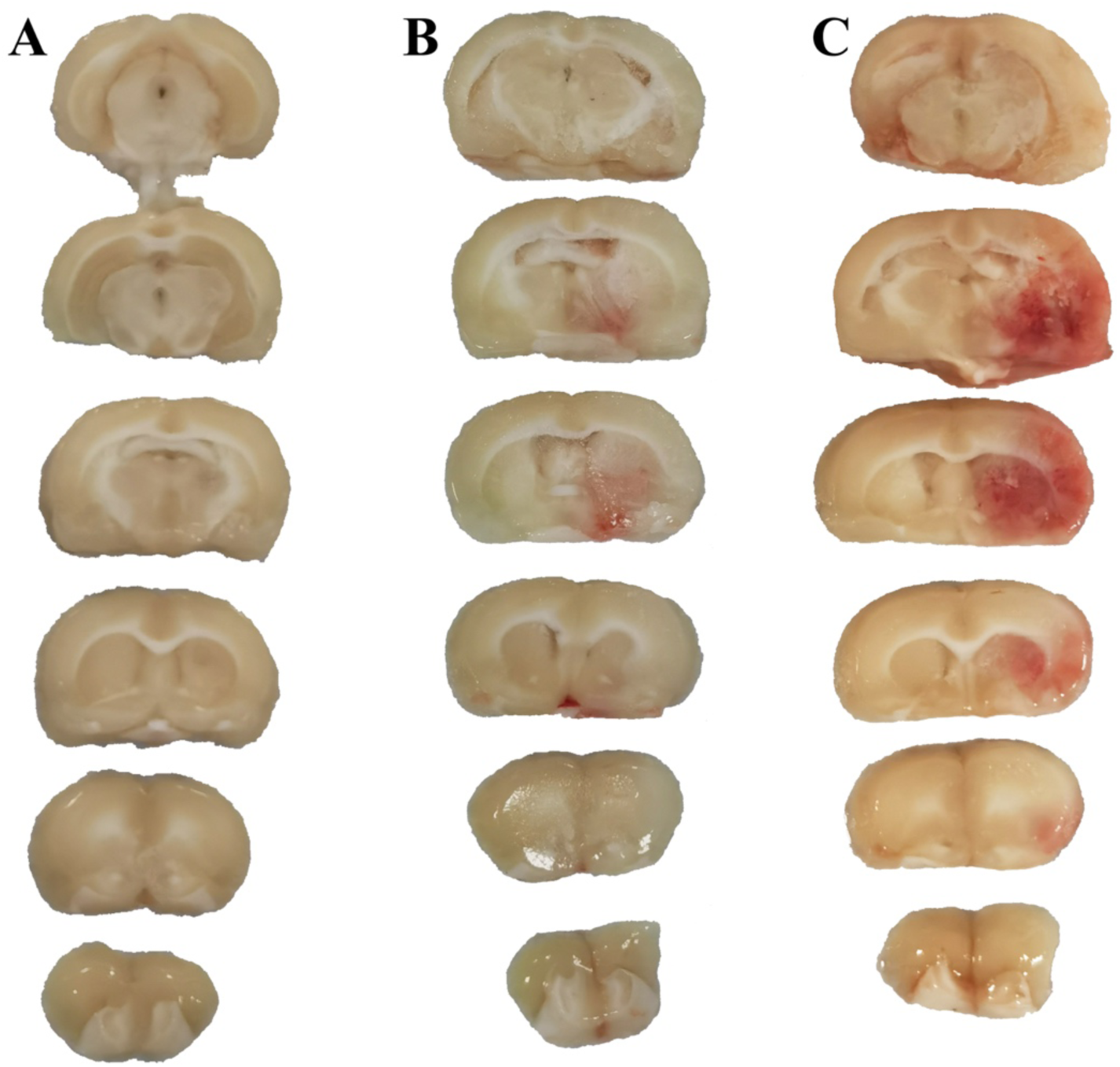
Macroscopic hemorrhagic transformation in the rat MCAO model was classified into no-hemorrhage (A), hemorrhagic infarction (B), and parenchymal hemorrhage (C).

### Primary cultures of neurons, astrocytes, microglia, BMECs, and pericytes

Sprague-Dawley rat neurons, astrocytes, microglia, BMECs, and pericytes were prepared as in previous studies [26–30]. Cortical primary neurons were isolated from the cerebral cortex of fetal rats on days 16-18 of pregnancy. In brief, the cortex was isolated, digested with 0.25% trypsin (Gibco) at 37 °C for 15 minutes, incubated with DNaseI stock solution for 30 seconds, and terminated the digestion with high sugar Dulbecco’s modified Eagle medium (DMEM) supplemented with 10% fetal bovine serum (FBS) complete medium (Gibco). The cell suspension was collected by a 100 µM nylon cell filter and centrifuged at 800 rpm for 5 minutes. Cells were resuspended with a complete medium and inoculated into the 0.01% Poly-L-lysine (Sigma) coated culture dish with 2.5x10^5^ cells per ml. Cells were placed in a humidified incubator with 5% CO_2_ at 37°C for 4 hours and further cultured with a complete medium containing Neurobasal, 2% B27, 2% GlutaMAX, and 1% Penicillin-Streptomycin (Gibco). Half of the culture medium was replaced every three days. Primary neurons were cultured for eight days.

Primary astrocytes were isolated from the cerebral cortex of newborn rats on days 1-2. The cortex was isolated, digested with 0.25% trypsin (Gibco) at 37 °C for 20 minutes, and terminated the digestion with a complete medium with DMEM-F12 (Gibco), 10% FBS, and 1% Penicillin-Streptomycin (Gibco). The cell suspension was collected by a 70 µM nylon cell filter and centrifuged at 1000 rpm for 5 minutes. Cells were resuspended with a complete medium and plated into the 75 cm^2^ flasks. Cells were cultured in a humidified incubator with 5% CO_2_ at 37°C with the complete medium replaced every 2-3 days.

Primary microglia were isolated from the cerebral cortex of newborn rats on days 1-2. The cortex was isolated, digested, and terminated the digestion, and the resulting cell suspension was collected by cell filter in the same way as astrocytes above. Then, the cell suspension was centrifuged at 1500 rpm for 5 minutes. Cells were resuspended with a complete medium containing DMEM-F12 (Gibco), 10% FBS, and 1% Penicillin-Streptomycin (Gibco) and plated into 0.01% Poly-L-lysine (Sigma) pre-coated 75 cm^2^ flasks. Cells were cultured in a humidified incubator with 5% CO_2_ at 37°C for two weeks, with the complete medium replaced every three days.

Subsequently, the loosely attached microglia were isolated by shaking the flasks for 4 hours at 200 rpm on a rotary shaker at 37°C. The cell suspension was then collected and centrifuged at 2000 rpm for 5 minutes. Cells were resuspended and cultured with a complete medium. Primary microglia were cultured for two days before treatment.

Primary BMECs were isolated from the cerebral cortex of rats on weeks 1-2. The cortex was isolated and digested with 0.1% type IV collagenase with 30 u/ml DNase I (Gibco) at 37 °C for 1.5 hours. The cell suspension was centrifuged at 1000 rpm for 8 minutes, suspended with 20% bovine serum albumin, and then centrifuged at 1000g for 20 minutes at 4°C. The sediment was digested with 0.1% collagenase/dispase enzyme containing 20 u/ml DNase I (Sigma) for 1 hour. The cell suspension was centrifuged at 1000 rpm for 8 minutes. Cells were suspended with DMEM, spread on 50% Percoll (Sigma) with a continuous gradient formed by centrifugation, and centrifuged at 25000g for 1 hour. The purified microvascular segments were collected, rinsed, and centrifuged at 1000 rpm for 5 minutes. Cells were resuspended with the endothelial cell medium (ECM) (ScienCell) and plated into the 25 cm^2^ flasks coated with type I collagen (Corning), and cultured in a humidified incubator with 5% CO_2_ at 37°C for 24 hours. Then cells were cultured with the ECM containing 4 ug/ml puromycin (Sigma) for 48 hours and replaced with the ECM without puromycin. After that, the ECM was changed every three days.

Primary pericytes were isolated from the cerebral cortex of rats on weeks 2-3. The cortex was isolated, minced, and homogenized in pre-cooled PBS, and then the homogenate was centrifuged at 500 rpm for 5 minutes at 4°C. The precipitation was suspended in 20% Percoll (Sigma) and centrifuged at 500 rpm for 20 minutes at 4°C. Next, the sediment was collected and digested with 0.1% collagenase II and 1000 U/mL DNase I (Sigma) at 37°C for 1 hour. After centrifugation at 500 rpm for 5 minutes at 4°C, the precipitation was harvested, resuspended in 20% Percoll, and centrifuged at 500 rpm for 15 minutes at 4°C. Subsequently, the middle layer was collected and centrifuged at 500 rpm for 5 minutes at 4°C. The sediment, isolated brain microvascular fragments containing pericytes and endothelial cells, was resuspended with the ECM for pericyte (ScienCell). Cells were cultured in a humidified incubator with 5% CO_2_ at 37°C for 24 hours. Then, cells were cultured with the ECM for pericyte containing 4 ug/ml puromycin (Sigma) for 48 hours and replaced with the ECM for pericyte without puromycin. After that, the ECM for pericyte was changed every three days. After 14 days of cultivation, excessive growth of pericytes and inhibition of endothelial cells were performed for passage purification.

The purity of primary neurons, astrocytes, microglia, BMECs, and pericytes were identified by immunofluorescence before subsequent experiments. The third-generation cells were used for experiments.

### OGD/R model

Rat neurons, astrocytes, microglia, BMECs, and pericyte cultures were separately exposed to OGD for a fixed time to mimic ischemia-like conditions in vitro [26–30]. Cell cultures were placed into a hypoxia chamber (Stemcell) at 37°C containing a gas mixture of 95% N2 and 5% CO_2_ and cultured in a glucose-free medium. Neurons, BMECs, pericytes, microglia, and astrocytes were separately maintained in the hypoxic chamber for 1 hour, 2 hours, 2 hours, 2 hours, and 6 hours. Then, cells were transferred into a cell incubator under normoxic conditions with a glucose-containing complete medium for reoxygenation mimicking reperfusion conditions. Cells were collected at the time of reoxygenation at 0, 3, 6, 12, 24, 48, and 72 hours. The cells in the control group did not receive OGD.

### RNA extraction and quantitative RT-PCR

The rat brains were quickly removed and dissected into the ischemic and non-ischemic hemispheres 24 hours after MCAO. Total RNA was extracted from the ischemic hemisphere’s peri-infarct and infarction core tissue using Trizol reagent (Invitrogen). Total RNA was extracted from the rat neurons, astrocytes, microglia, BMECs, and pericytes after OGD/R. Good RNA quality was considered an OD260/OD280 ratio between 1.8 and 2.0.

Twenty-nine differentially expressed microRNAs in the ischemic hemispheres from the rat reperfusion-induced HT model examined by the microRNA array were preliminarily shown in our previous study [23]. In this study, twenty-five differentially expressed microRNAs shown from the microRNA array (except four microRNAs not expressed in the human and mouse) were assessed by quantitative RT-PCR between PH group and sham-operated group in the MCAO model (n=5, each group). Then, significantly differentially expressed PH-related microRNAs confirmed in the peri-infarct of ischemic hemisphere in the MCAO model were further assessed in the OGD/R model of neurons, astrocytes, microglia, BMECs, and pericytes.

The expression of mature microRNAs and mRNA of predicted target genes from the later bioinformatics analysis in the peri-infarct and infarction core tissue and cells after OGD/R were assessed using the TaqMan stem-loop and the SYBR green real-time RT-PCR method. We performed quantitative RT-PCR using the PCR Master Mix on the Applied Biosystem PCR System. MicroRNAs and mRNA expression were normalized using U6 and β-actin as an internal control and were calculated using the 2-ΔΔCT method. The primer sequences for microRNAs and predicted target mRNAs are shown in Table S1 and Table 1.

**Table 1.**
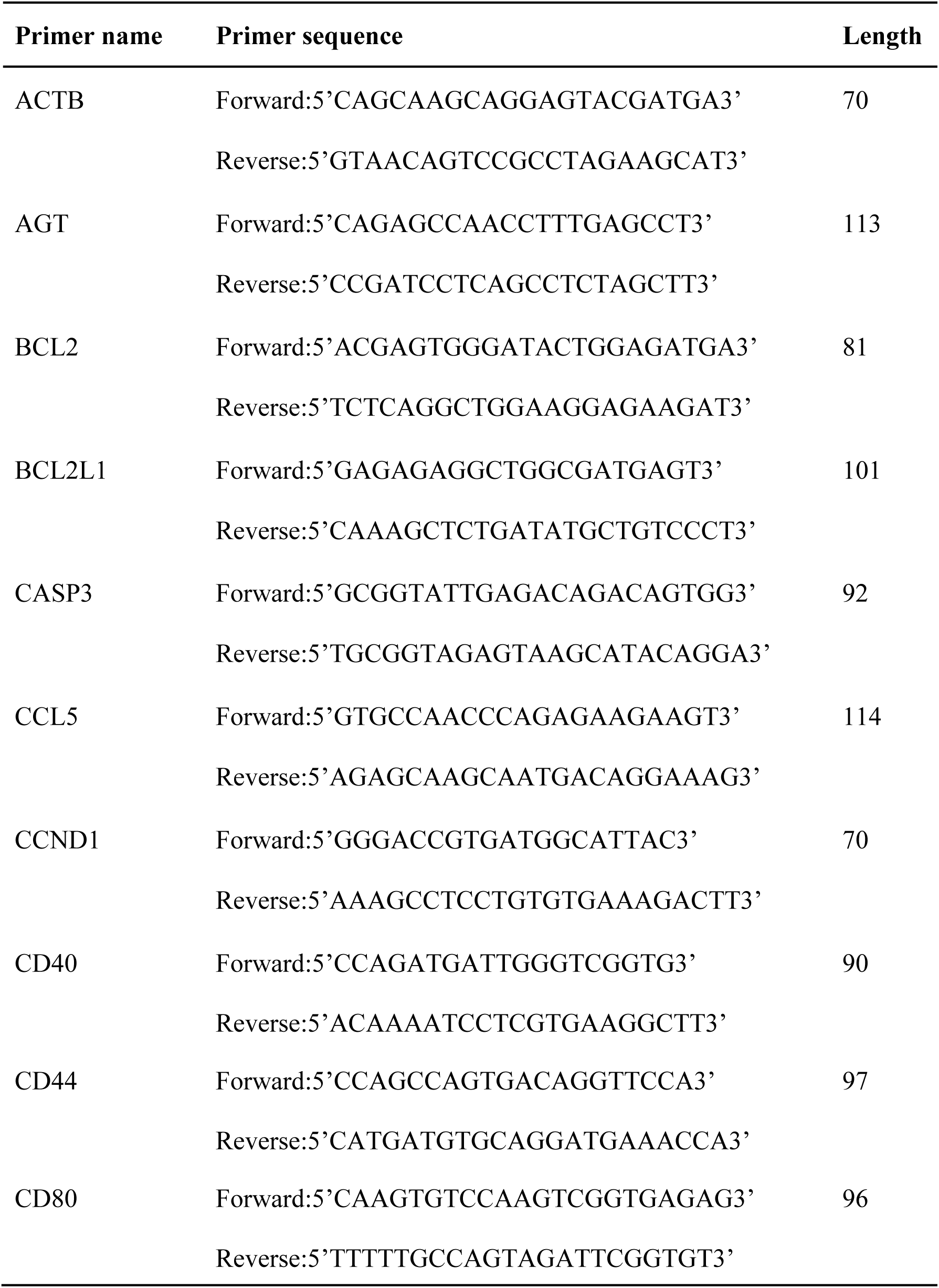
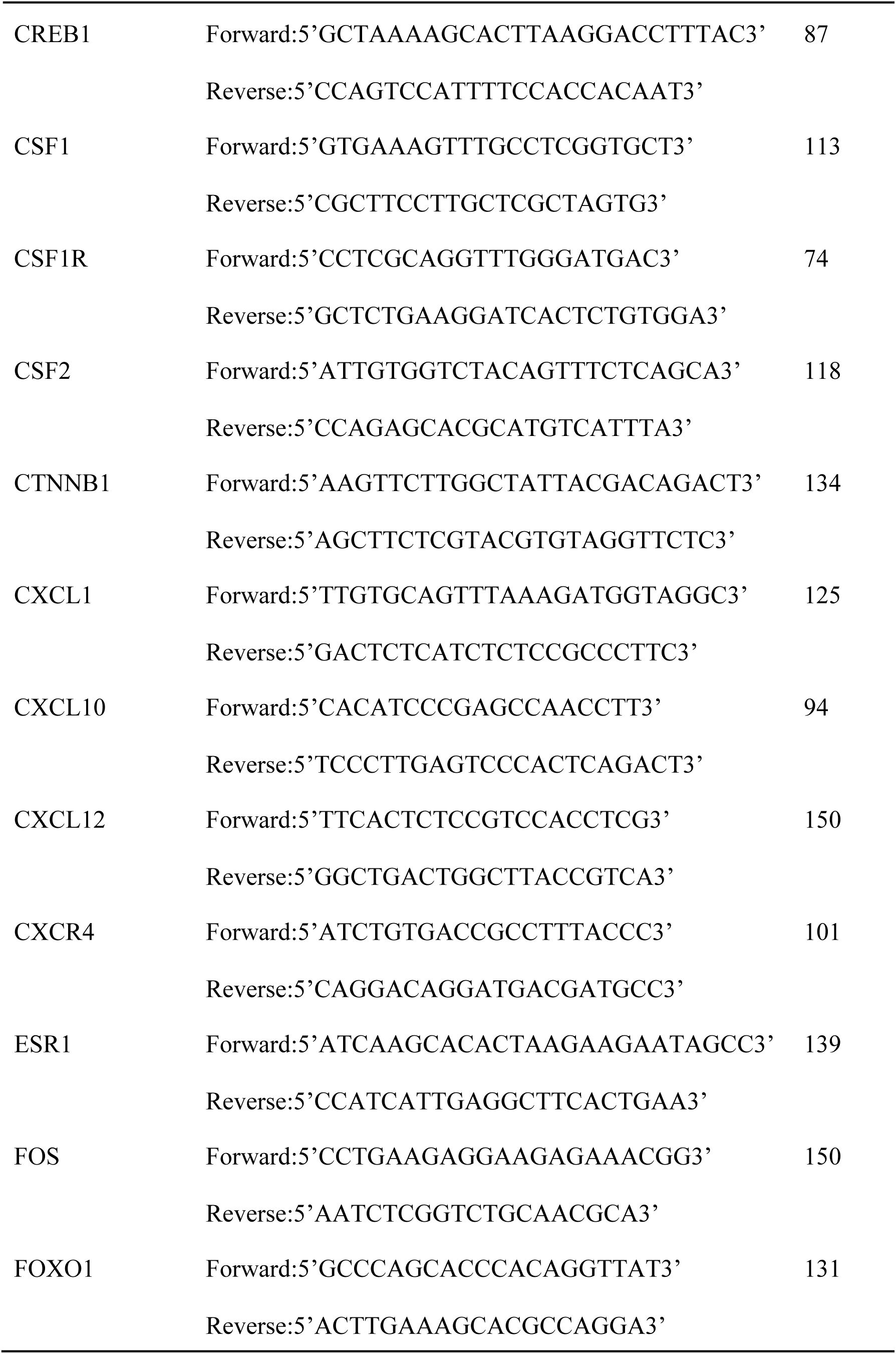
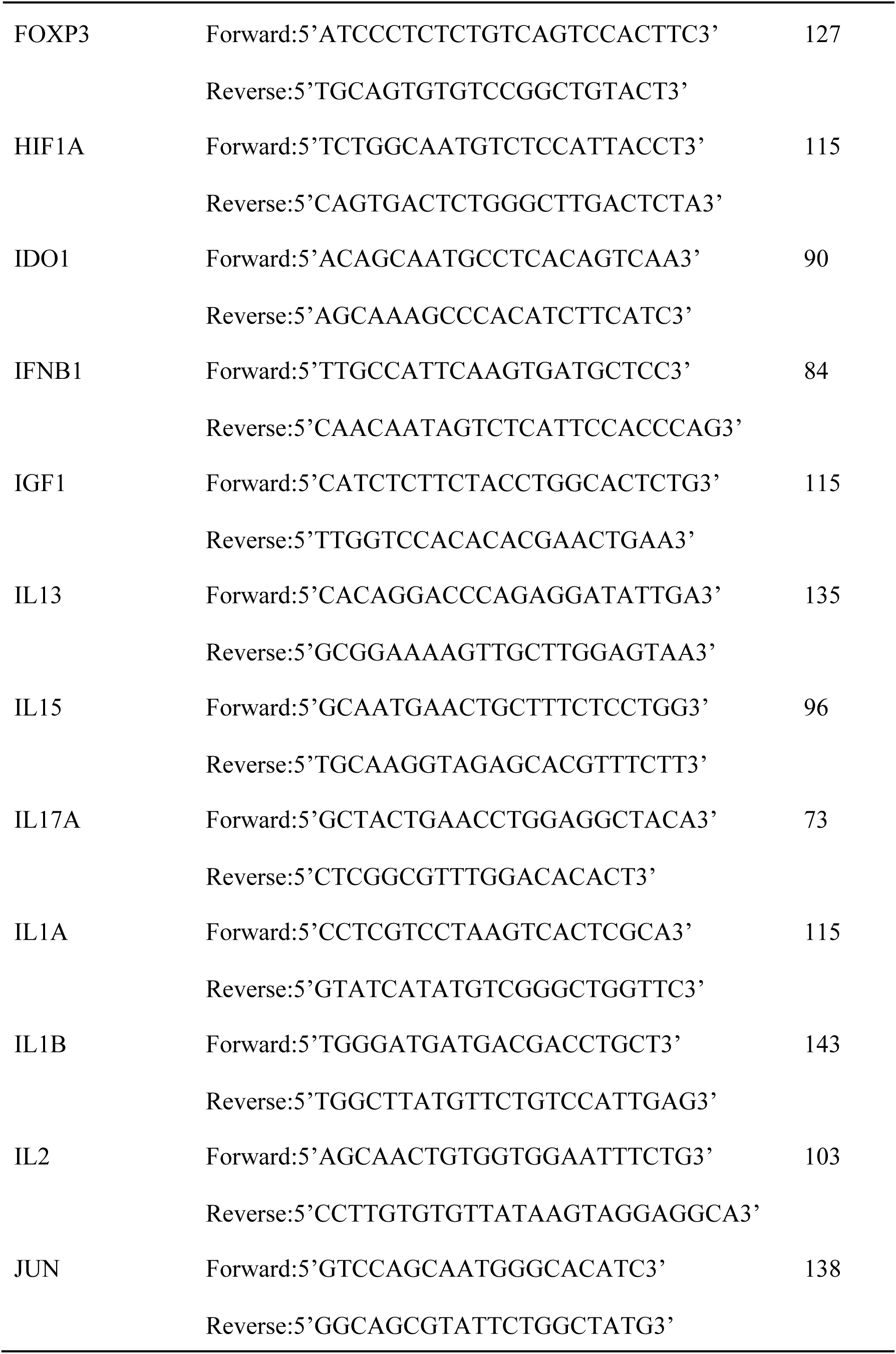
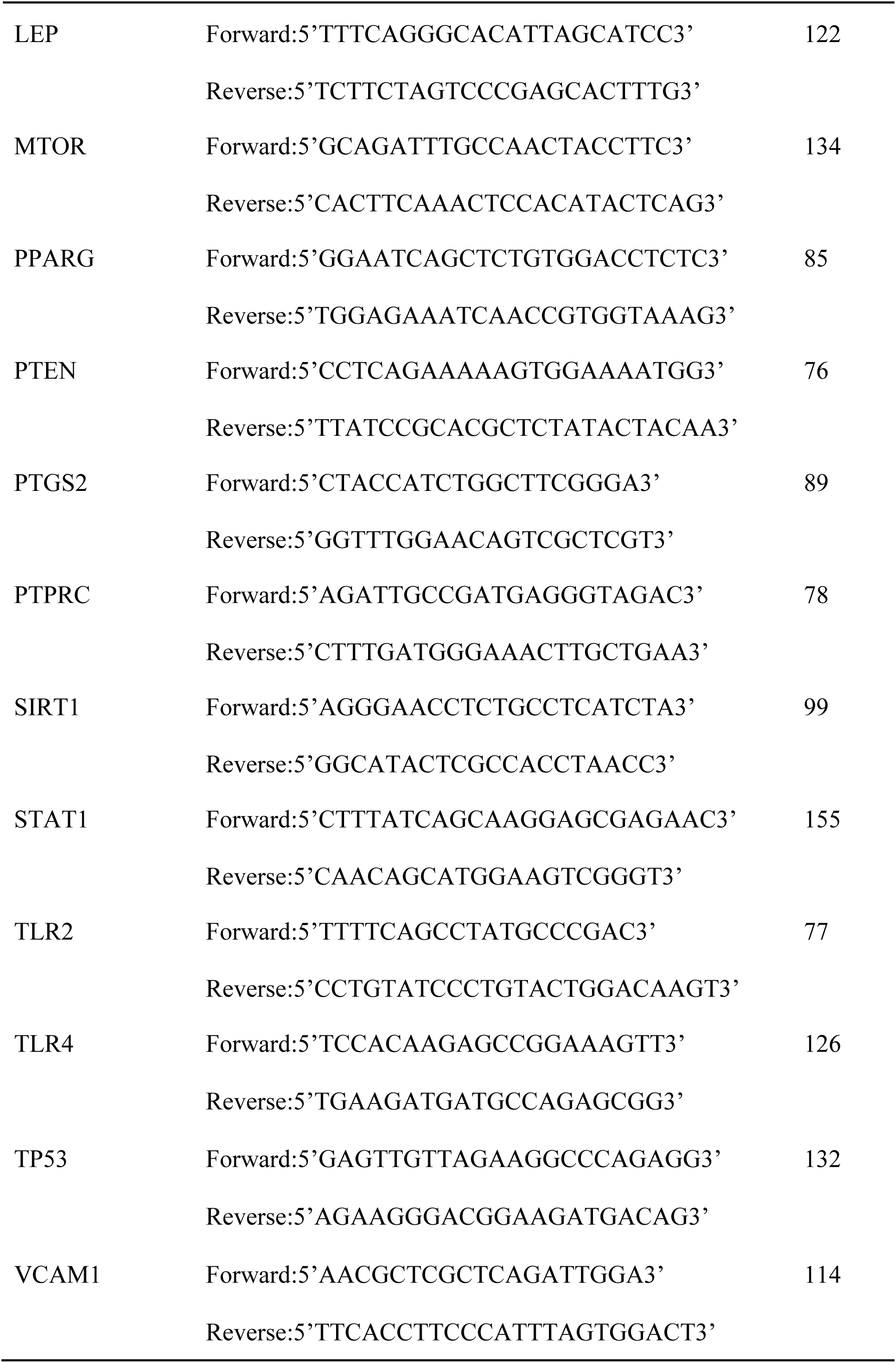
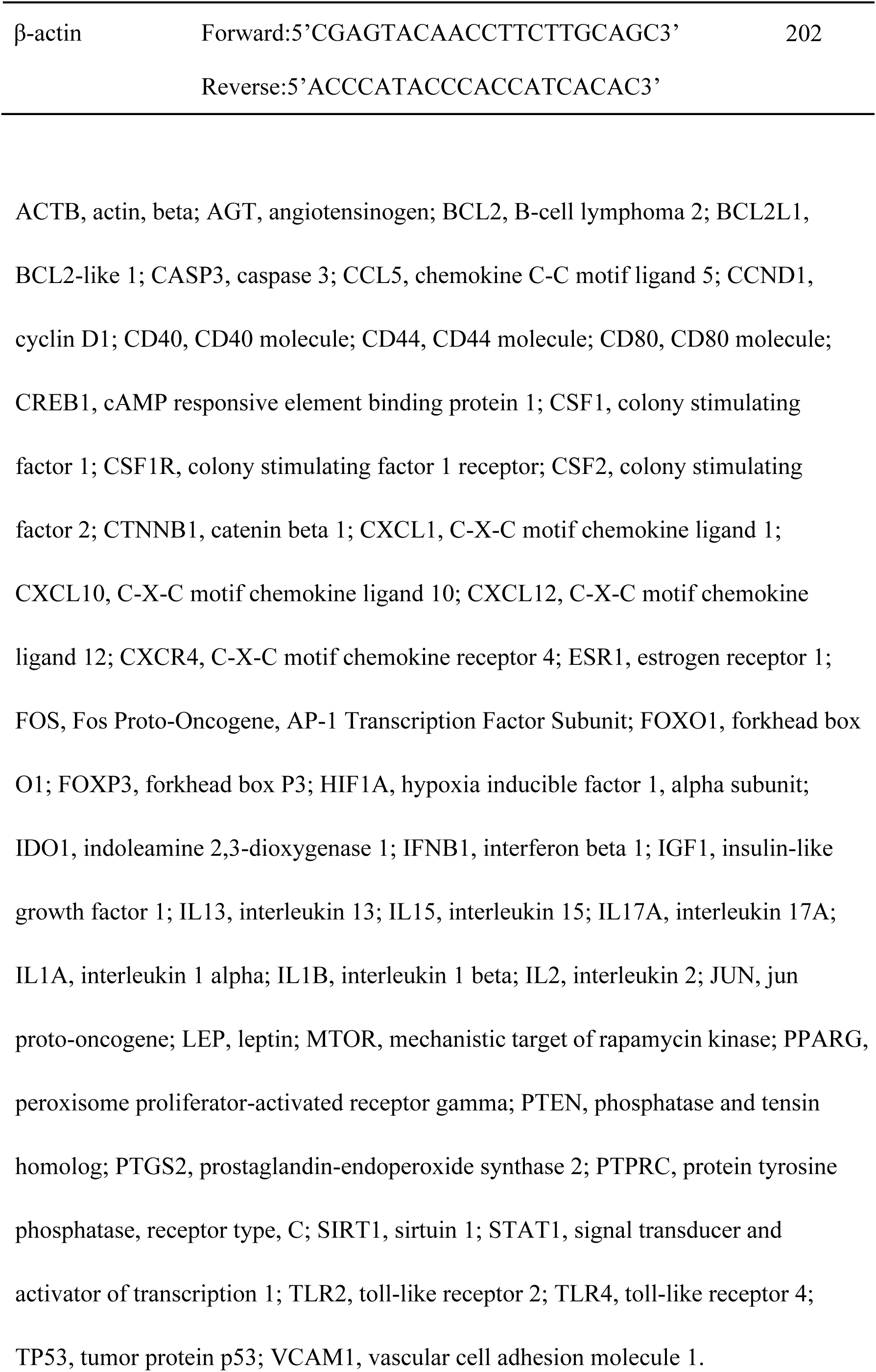
Sequences of the primer for key predicted target mRNAs used in the qRT-PCR.

### Gene Ontology (GO) and Kyoto Encyclopedia of Genes and Genomes (KEGG)

#### Pathway Analysis

The significantly differentially expressed PH-related microRNAs confirmed in the peri-infarct of the ischemic hemisphere in the rat MCAO model were selected for the bioinformatics analysis and target gene prediction. We searched the predicted target genes from differentially expressed microRNAs by TargetScan or miRDB databases. KEGG pathway and GO functional enrichment analysis were carried out to reveal the biological function of the predicted target genes using the cluster Profiler R package (https://yulab-smu.top/biomedical-knowledge-mining-book/; version 4.11.0) [31]. The biological process, cellular component, and molecular function of the predicted target genes were revealed from the GO enrichment analysis. KEGG and GO terms with an adjusted P value less than 0.05 were considered significantly enriched.

### Construction of microRNA-mRNA regulatory network of PH

We obtained the immune-related genes, inflammation-related genes, oxidative stress-related genes, and apoptosis-related genes from the GeneCards database (http://www.genecards.org; version 5.18). The Search Tool for the Retrieval of Interacting Genes (STRING) (https://cn.string-db.org/cgi/input; version 11.5) was applied to identify the protein-protein interaction network of the predicted target genes related to the mechanism of immune, inflammation, oxidative stress, and apoptosis. The top twenty hub genes associated with immune, inflammation, oxidative stress, and apoptosis mechanisms were shown using the maximal clique centrality algorithm in the CytoHubba, a plugin in Cytoscape software. Then, the predicted microRNA-mRNA network containing hub genes was established by the online tool Cytoscape (https://cytoscape.org; version 3.9.1). Finally, we obtained the microRNA-mRNA regulatory network of PH after the expression of the predicted hub genes was confirmed by RT-PCR in the rat reperfusion-induced PH model.

### Statistical analysis

We performed the statistical analyses using SPSS 22.0 software. The continuous and categorical variables were analyzed by Student t-test, Mann-Whitney U test, and one-way ANOVA. A p-value less than 0.05 was considered significant.

## Results

### MicroRNAs in rat ischemic brain with PH after mechanical reperfusion

HT was shown in all 42 rats from the transient MCAO model. The percentage of PH was 43% in the transient MCAO group. This reperfusion-induced hemorrhage model had a 57% mortality, mainly within 12 hours after reperfusion. The survival of ten rats in the transient MCAO group with PH and ten in the sham-operated group was further used. Twenty-five differentially expressed microRNAs shown from our previous microRNA array study were further confirmed by quantitative RT-PCR. Finally, 14 microRNAs were significantly differentially expressed in both peri-infarct and infarction core in the rats with PH compared with the sham-operated rats, including 13 down-regulated microRNAs and one up-regulated microRNA (miRNA-195-3p). These 13 down-regulated microRNAs associated with PH included miRNA-126a-5p, miRNA-132-3p, miRNA-136-3p, miRNA-142-3p, miRNA-153-5p, miRNA-218a-5p, miRNA-219a-2-3p, miRNA-29a-5p, miRNA-29c-3p, miRNA-369-5p, miRNA-376a-5p, miRNA-376b-5p, and miRNA-383-5p (Figure 2 and Table S2).

**Figure 2.**
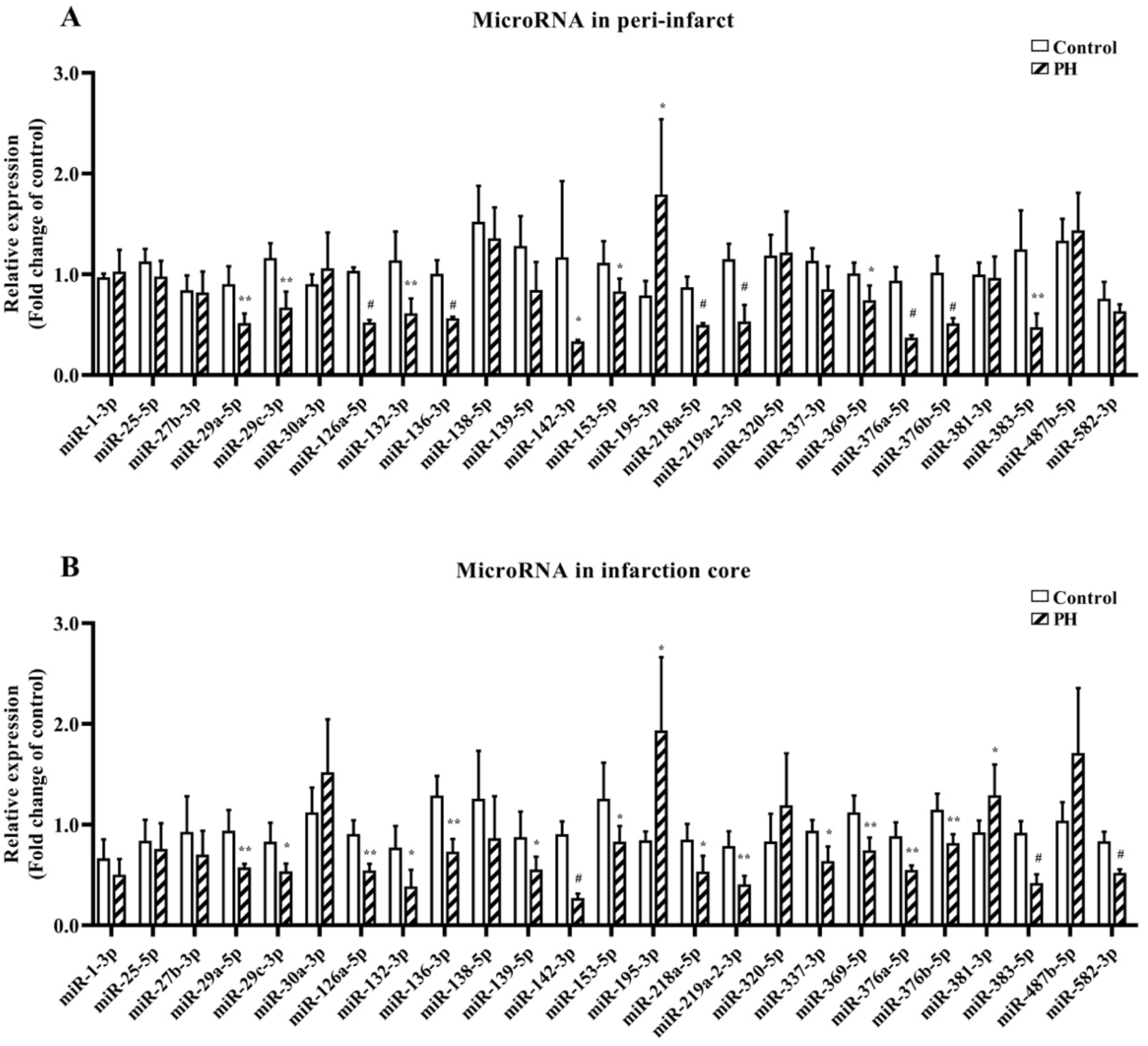
Twenty-five microRNAs shown from our previous microRNA array study were confirmed in the peri-infarct (A) and infarction core (B) by quantitative RT-PCR. Fourteen microRNAs were significantly differentially expressed in the rats with parenchymal hemorrhage compared with the sham-operated rats, including 13 down-regulated microRNAs and one up-regulated microRNA. *P<0.05; **P<0.01; #P<0.001.

### Differential expression of PH-related microRNAs in the OGD/R model

These 14 significantly differentially expressed microRNAs associated with PH from the rat MCAO model were further assessed in the OGD/R model with neurons, astrocytes, microglia, BMECs, and pericytes (Table S3 and Figure S1-S5). We identified 11 microRNAs with a significantly different abundance level at 24 hours in neurons treated by OGD/R compared with the non-OGD/R control. Eight down-regulated microRNAs were consistent with the results of the animal model, including miRNA-132-3p, miRNA-136-3p, miRNA-153-5p, miRNA-218a-5p, miRNA-369-5p, miRNA-376a-5p, miRNA-376b-5p, and miRNA-383-5p (Figure 3A).

**Figure 3.**
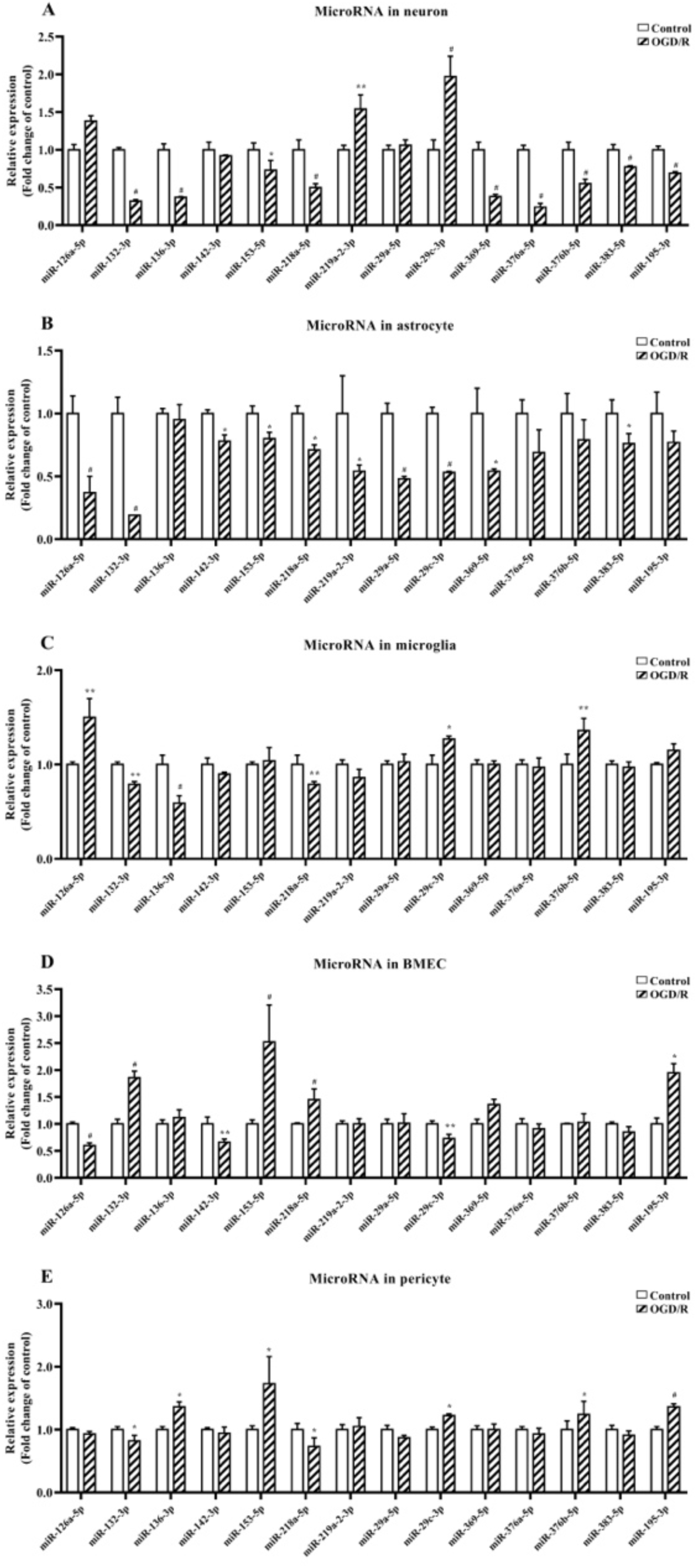
Differential expression of 14 PH-associated microRNAs in neuron, astrocyte, microglia, brain microvascular endothelial cells (BMEC), and pericyte after oxygen-glucose deprivation/reoxygenation (OGD/R) at 24 hours. A, microRNAs in the OGD/R model of neuron. B, microRNAs in the OGD/R model of astrocyte. C, microRNAs in the OGD/R model of microglia. D, microRNAs in the OGD/R model of BMEC. E, microRNAs in the OGD/R model of pericyte. *P<0.05; **P<0.01; #P<0.001.

We identified ten microRNAs that had significantly different abundance levels at 24 hours in the OGD/R-treated astrocytes compared with non-OGD/R control, all of which were decreased and consistent with the animal model results. These down-regulated 10 microRNAs included miRNA-126a-5p, miRNA-132-3p, miRNA-142-3p, miRNA-153-5p, miRNA-218a-5p, miRNA-219a-2-3p, miRNA-29a-5p, miRNA-29c-3p, miRNA-369-5p, and miRNA-383-5p (Figure 3B). Six microRNAs had a significantly different abundance level at 24 hours in the OGD/R-treated microglia compared with non-OGD/R control. Three down-regulated microRNAs were consistent with the results of the animal model, including miRNA-132-3p, miRNA-136-3p, and miRNA-218a-5p (Figure 3C).

We identified seven microRNAs with a significantly different abundance level at 24 hours in the OGD/R-treated BMECs compared with non-OGD/R control. Two down-regulated microRNAs (miRNA-126a-5p and miRNA-142-3p) and one up-regulated microRNA (miRNA-195-3p) were consistent with the results of the MCAO model (Figure 3D). Seven microRNAs had a significantly different abundance level at 24 hours in the OGD/R-treated pericytes compared with non-OGD/R control. Two down-regulated microRNAs (miRNA-132-3p and miRNA-218a-5p) and one up-regulated microRNA (miRNA-195-3p) were consistent with the results of the MCAO model (Figure 3E).

Ten microRNAs were significantly differentially expressed in at least two of five models of neuron, astrocyte, microglia, BMEC, and pericyte after OGD/R at 24 hours, consistent with the results of the animal model. Both miRNA-132-3p and miRNA-218a-5p significantly decreased in the model of neurons, astrocytes, microglia, and pericytes after OGD/R at 24 hours. MiRNA-126a-5p, miRNA-142-3p, and miRNA-29c-3p significantly decreased in the model of astrocytes and BMEC after OGD/R at 24 hours. MiRNA-153-5p, miRNA-369-5p, and miRNA-383-5p significantly decreased in the model of neurons and astrocytes after OGD/R at 24 hours. MiRNA-136-3p significantly decreased in the model of neurons and microglia after OGD/R at 24 hours. MiRNA-195-3p significantly increased in the model of BMEC and pericyte after OGD/R at 24 hours.

### KEGG pathway and GO analysis of differentially expressed microRNAs associated with PH

To identify the potential biological function of these 14 differentially expressed microRNAs, we found 9992 target genes for these microRNAs predicted by TargetScan or miRDB databases (Table S4). Then, KEGG and GO enrichment analyses were performed. The top ten enriched terms of KEGG pathways, biological process, cellular component, and molecular function of GO are archived (Figure 4 and Table S5). The top ten enriched pathways were shown as follows: MAPK signaling pathway, Ras signaling pathway, Rap1 signaling pathway, regulation of actin cytoskeleton, proteoglycans in cancer, Wnt signaling pathway, signaling pathways regulating pluripotency of stem cells, breast cancer, FoxO signaling pathway, growth hormone synthesis, secretion and action.

**Figure 4.**
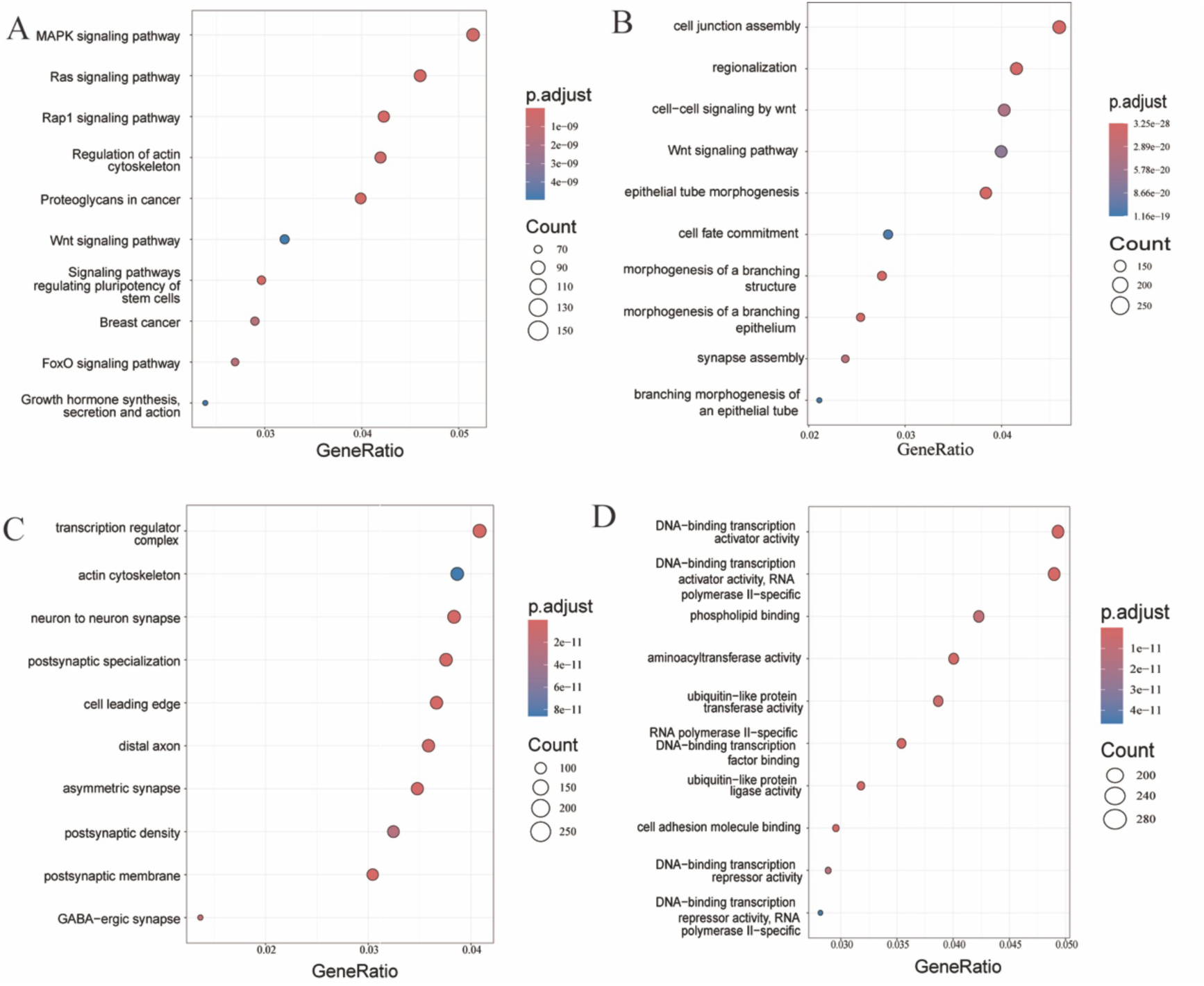
Kyoto Encyclopedia of Genes and Genomes (KEGG) pathway and Gene Ontology (GO) functional enrichment analysis of the predicted target genes from differentially expressed microRNAs. (A) KEGG. (B) GO-biological processes. (C) GO-cell components. (D) GO-molecular functions.

### Differential expression of predicted target genes in the rat PH model

The immune-related genes, inflammation-related genes, oxidative stress-related genes, and apoptosis-related genes were obtained from the GeneCards database. There were six hundred and thirty-three, five hundred and seven, five hundred and sixty-five, and eight hundred and eleven predicted target genes related to immune, inflammation, oxidative stress, and apoptosis, respectively (Table S6). Using the STRING online tool, we identified the protein-protein interaction network of these predicted target genes related to immune, inflammation, oxidative stress, and apoptosis. Then, the top twenty hub genes related to immune, inflammation, oxidative stress, and apoptosis were calculated in the CytoHubba plugin, respectively (Figure 5A-D and Table S7). Forty-six key predicted target genes for these 14 PH-related microRNAs were identified since 28 genes involved at least two mechanisms with immune, inflammation, oxidative stress, or apoptosis (Table S8). Then, we predicted the microRNA-mRNA network of PH, including 107 microRNA-mRNA regulatory axes containing 46 hub genes using the Cytoscape software (Figure 5E).

**Figure 5.**
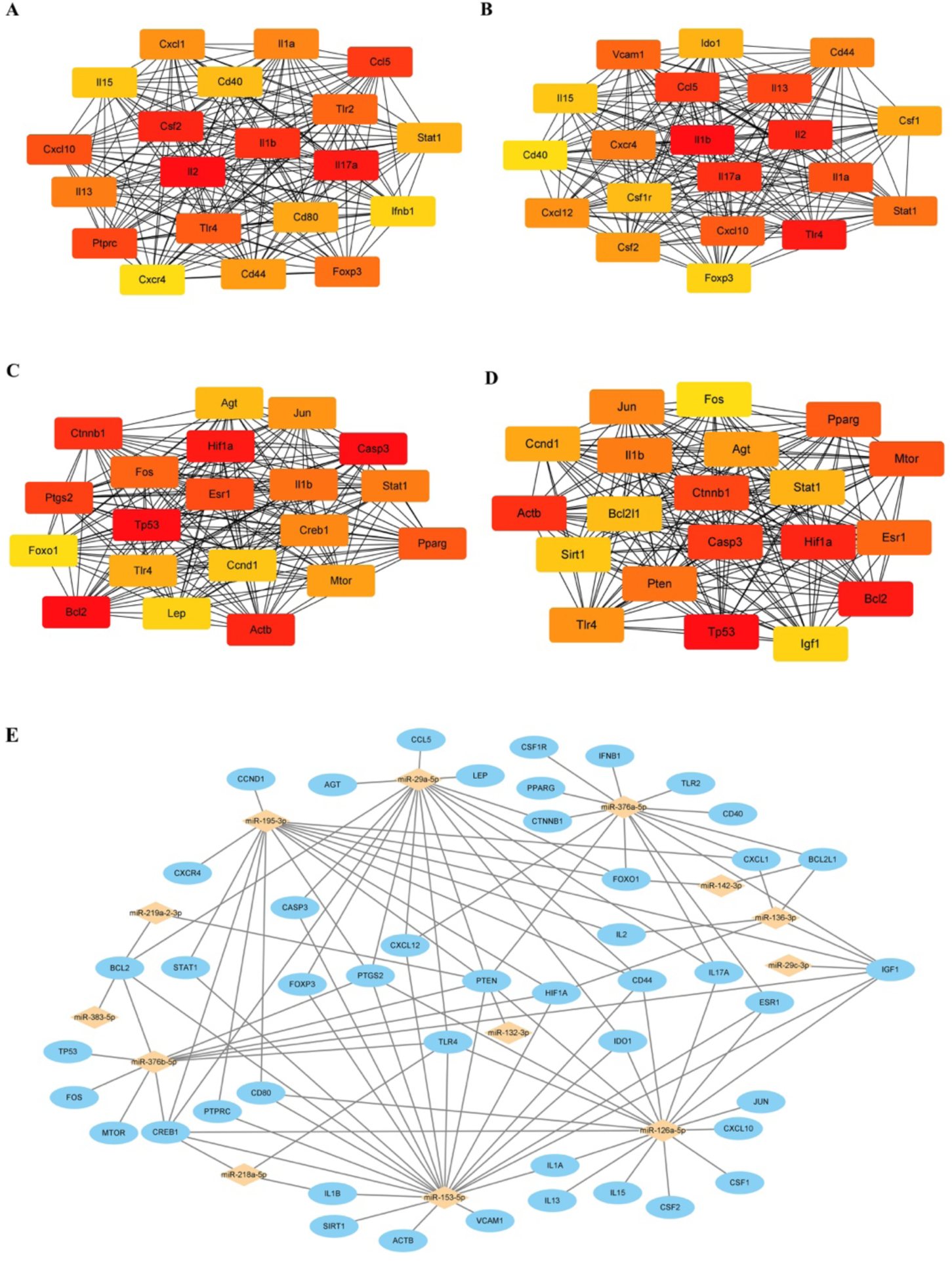
The predicted microRNA-mRNA regulatory network of PH. Relationship network diagram of the top twenty hub genes related to immune (A), inflammation (B), oxidative stress (C), and apoptosis (D) from the protein-protein interaction network. (E) The reconstructed 107 microRNA-mRNA regulatory axes with 46 predicted hub genes.

These 46 predicted hub genes were further assessed by RT-PCR in the animal model. Thirty-one genes were significantly differentially expressed in peri-infarct tissue in rats with PH compared to sham-operated rats. Twenty-two up-regulated genes associated with PH were confirmed, including caspase 3 (CASP3), chemokine C-C motif ligand 5 (CCL5), cyclin D1 (CCND1), CD44 molecule (CD44), CD80, CSF (colony stimulating factor)1, CSF2, CXCL (C-X-C motif chemokine ligand)1, CXCL10, ESR (estrogen receptor)1, Fos Proto-Oncogene, AP-1 Transcription Factor Subunit (FOS), hypoxia inducible factor 1 alpha subunit (HIF1A), indoleamine 2,3-dioxygenase 1 (IDO1), interferon beta 1 (IFNB1), insulin-like growth factor 1 (IGF1), insulin-like growth factor 1 (IL1A), interleukin 1 beta (IL1B), jun proto-oncogene (JUN), prostaglandin-endoperoxide synthase 2 (PTGS2), protein tyrosine phosphatase receptor type C (PTPRC), toll-like receptor (TLR)2, and TLR4. Nine down-regulated genes associated with PH included angiotensinogen (AGT), BCL2-like 1 (BCL2L1), cAMP responsive element binding protein 1 (CREB1), catenin beta 1 (CTNNB1), CXCL12, forkhead box O1 (FOXO1), mechanistic target of rapamycin kinase (MTOR), signal transducer and activator of transcription 1 (STAT1), and vascular cell adhesion molecule 1 (VCAM1) (Figure 6 and Table S9). Finally, we confirmed twenty-five, twenty-two, twenty, and twenty-five PH-associated genes involved in the mechanism of immune, inflammation, oxidative stress, and apoptosis, respectively. Eight up-regulated genes (CCL5, FOS, HIF1A, IL1A, IL1B, JUN, PTGS2, TLR4) and four down-regulated genes (CREB1, CTNNB1, MTOR, STAT1) were involved in all of immune, inflammation, oxidative stress, and apoptosis.

**Figure 6.**
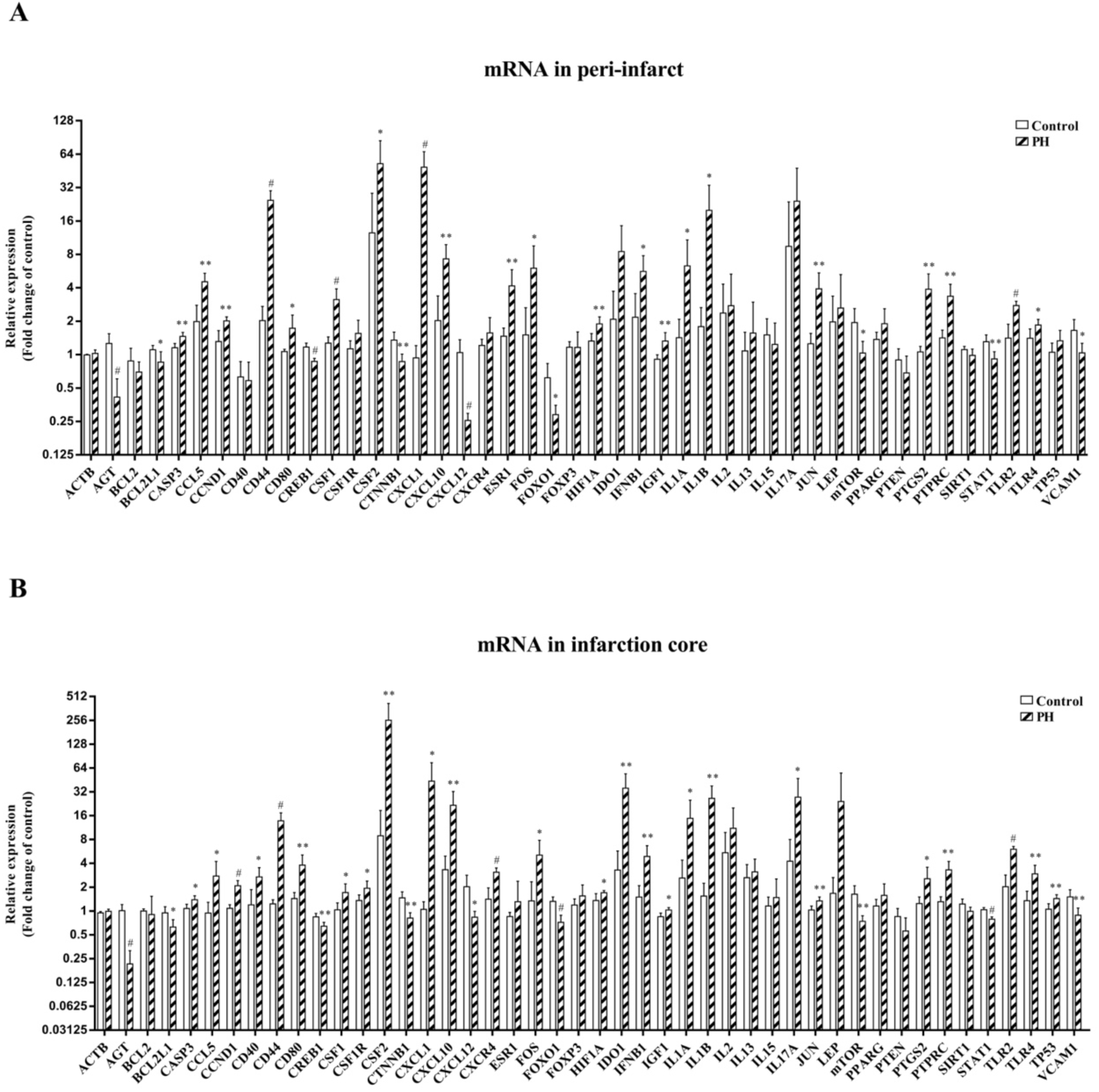
Forty-six predicted hub genes from differentially expressed microRNAs were assessed by quantitative RT-PCR in the peri-infarct (A) and infarction core (B). Thirty-one genes were significantly differentially expressed in peri-infarct tissue in rats with parenchymal hemorrhage compared to sham-operated rats, including 22 up-regulated genes and nine down-regulated genes. *P<0.05; **P<0.01; #P<0.001.

### The microRNA-mRNA regulatory network of PH

Finally, we obtained the microRNA-mRNA network of PH based on the prediction tool and PCR examination, including nine microRNAs and their 24 essential target genes with 49 microRNA-mRNA regulatory axes (Figure 7 and Table S10). The microRNA-mRNA regulatory network of PH was associated with the mechanism of immune, inflammation, oxidative stress, and apoptosis. For example, we found decreased miRNA-218a-5p and increased expression of its two key predicted target genes (IL1B and TLR4) in the peri-infarct tissue in the rat model with PH. In the OGD/R model of neurons, astrocytes, microglia, and pericytes, the expression of miRNA-218a-5p significantly decreased with the increased expression of the predicted target gene of IL1B and TLR4 (Figure 8). In addition, miRNA-195-3p was up-regulated, and its three key predicted target genes (CREB1, FOXO1, STAT1) were decreased in the peri-infarct tissue. In the OGD/R model of BMEC and pericyte, miRNA-195-3p significantly increased with the decreased expression of the predicted target gene of FOXO1 and STAT1 (Figure 9).

**Figure 7.**
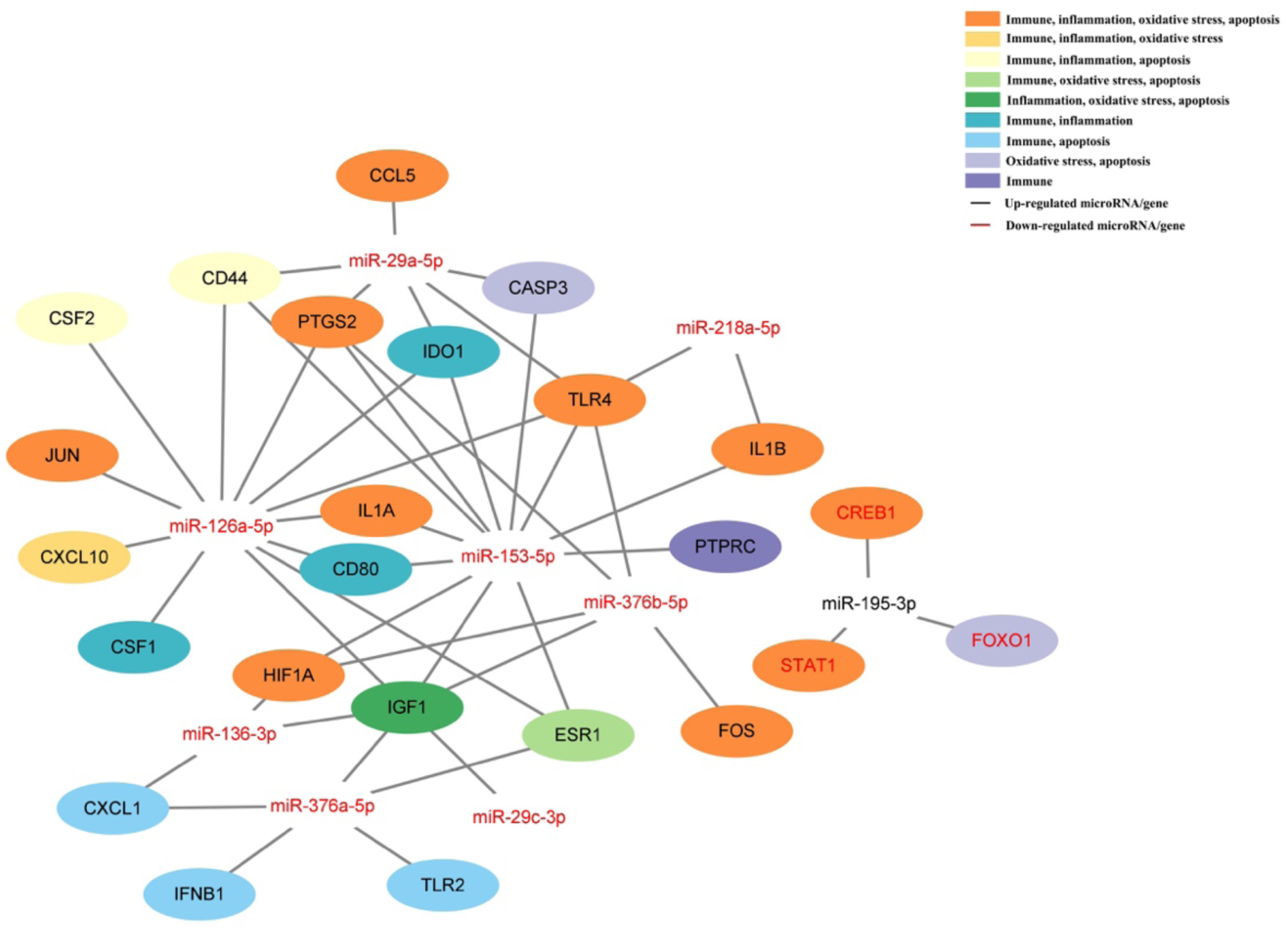
The microRNA-mRNA regulatory axes of parenchymal hematoma based on the prediction tool and PCR examination, including nine microRNAs (eight down-regulated microRNAs and one up-regulated microRNA) and their 24 essential target genes (three down-regulated genes and 21 up-regulated genes) with 49 microRNA-mRNA regulatory axes. This network was associated with the mechanism of immune, inflammation, oxidative stress, and apoptosis.

**Figure 8.**
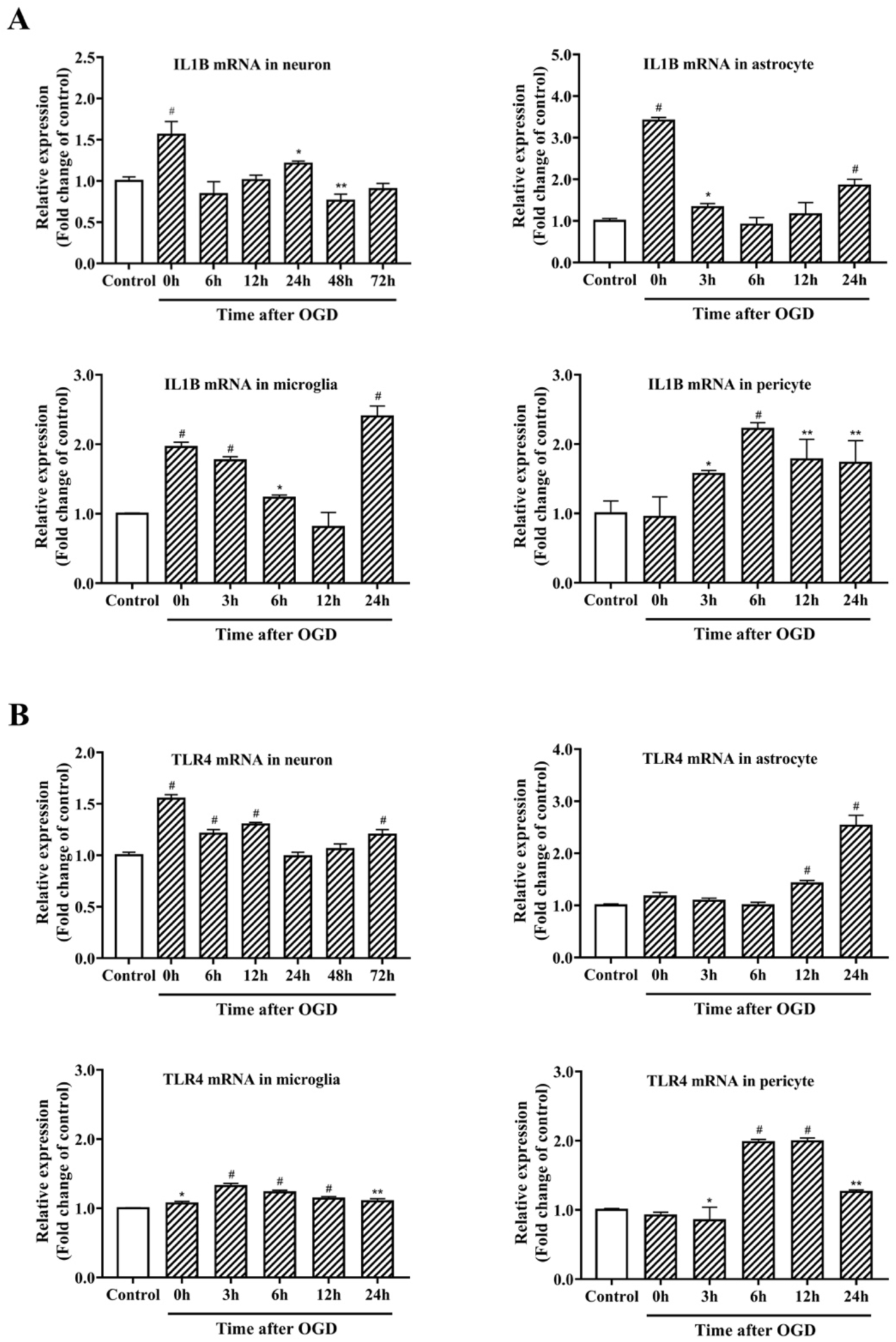
The expression of interleukin 1B (IL1B) and toll-like receptor 4 (TLR4) as the predicted target gene of microRNA-218a-5p increased in the oxygen-glucose deprivation/reoxygenation model of neuron, astrocyte, microglia, and pericyte. (A) IL1B. (B) TLR4.

**Figure 9.**
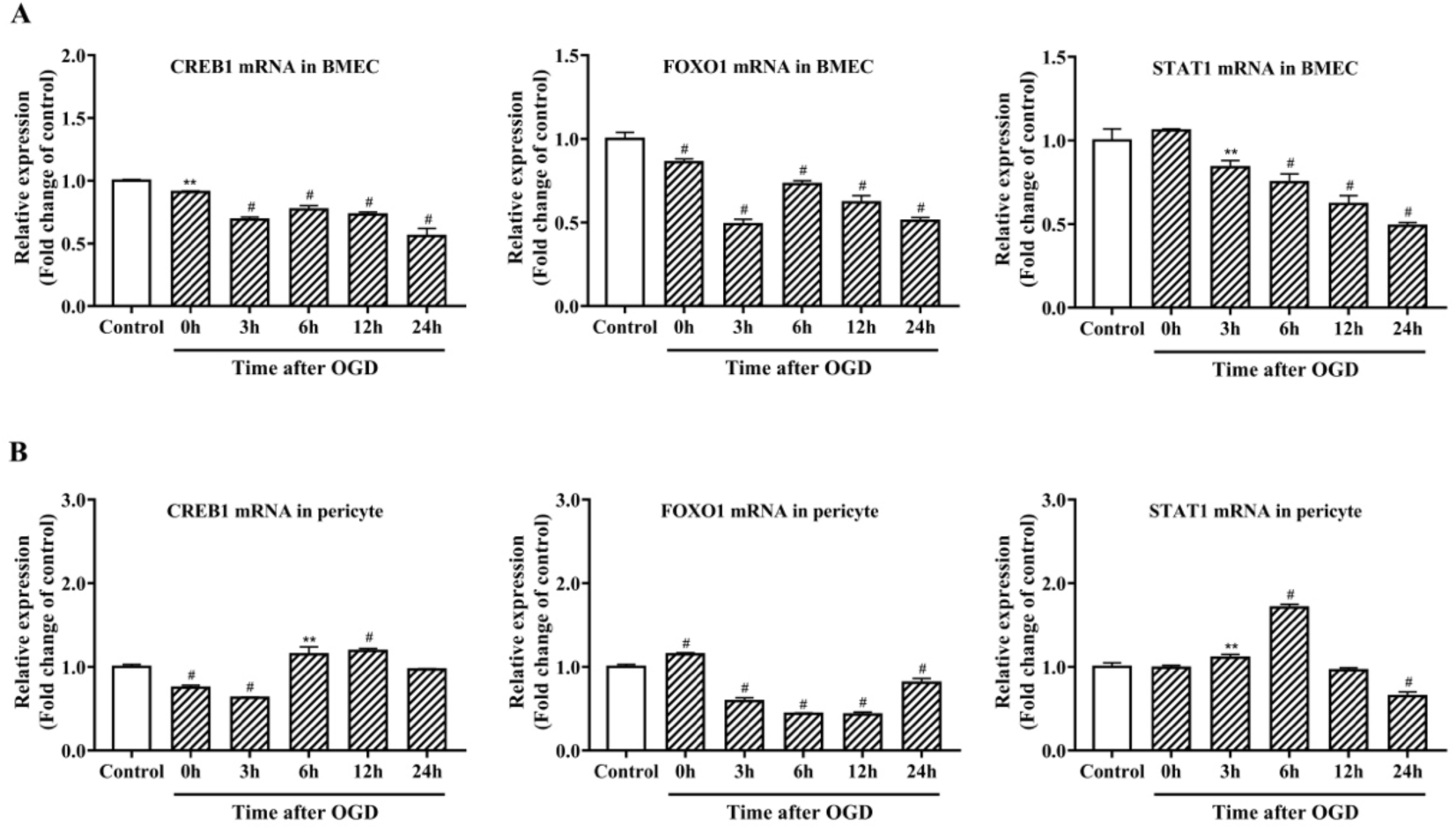
The expression of cAMP responsive element binding protein 1 (CREB1), forkhead box O1 (FOXO1), and signal transducer and activator of transcription 1 (STAT1) as the predicted target gene of miR-195-3p was shown in the oxygen-glucose deprivation/reoxygenation model of brain microvascular endothelial cell (BMEC) and pericyte. (A) genes in the model of BMEC. (B) genes in the model of pericyte. *P<0.05; **P<0.01; #P<0.001.

## Discussion

This study showed 13 down-regulated and one up-regulated microRNAs in both peri-infarct and infarction core with PH from the hyperglycemic rat MCAO model after mechanical reperfusion. Among these 14 PH-related microRNAs, ten microRNAs were significantly differentially expressed in at least two of five models of neuron, astrocyte, microglia, BMEC, and pericyte after OGD/R. MAPK signaling pathway, Ras signaling pathway, Rap1 signaling pathway, and Wnt signaling pathway were the enriched pathway of these PH-related microRNAs. MicroRNAs contribute to cerebral ischemic injury by regulating oxidative stress, inflammation, apoptosis, and BBB permeability [22, 32–34]. In this study, we showed 31 significantly differentially expressed target genes related to immune, inflammation, oxidative stress, and apoptosis which were predicted from these PH-associated microRNAs. Forty-nine microRNA-mRNA regulatory axes of PH were revealed.

Eight microRNAs from these 14 PH-related microRNAs have been reported as a potential regulator for cerebral ischemia or reperfusion injury in previous studies [19, 22, 32–34]. For example, our study showed that miRNA-126-5p decreased in the peri-infarct of rat MCAO model with PH and in BMECs after OGD/R at 24 hours. In a recent study using the mouse MCAO model, miRNA-126-5p significantly decreased in the brain tissue. Overexpression of the miRNA-126-5p attenuated BBB disruption after ischemic stroke [35]. Our study showed the other six differentially expressed microRNAs associated with PH may be candidates to ameliorate cerebral ischemia, including miRNA-29a-5p, miRNA-219a-2-3p, miRNA-369-5p, miRNA-376a-5p, miRNA-376b-5p, and miRNA-195-3p.

Our study provides new evidence for the association of microRNAs with PH in AIS after mechanical reperfusion therapy. Most differentially expressed microRNAs were not reported to be involved in PH after reperfusion [20, 21, 36–41].

Overexpression of miRNA-21-5p, miRNA-206, and miRNA-3123 were associated with HT in patients with cardioembolic stroke. These three microRNAs related to matrix metalloproteinase-9 may serve as prognostic blood markers for HT in cardioembolic stroke [20, 36]. A recent study revealed the expression levels of non-coding RNA in peripheral blood mononuclear cells associated with HT in six patients with AIS receiving EVT. There were 47 significantly differentially expressed microRNAs from peripheral blood in patients with HT compared with those without HT. MiRNA-383-5p was one of the top ten down-regulated microRNAs related to HT, similar to our study [21]. In the transient MCAO model of wild-type mice, miRNA-155 increased in the peri-infarct regions, which regulated BBB permeability via direct interactions with several mRNAs. Genetic deletion of miRNA-155 significantly reduced the hemorrhagic burden after cerebral ischemic reperfusion [37]. In addition, we found 10 PH-related microRNAs significantly differentially expressed in at least two of five models of neuron, astrocyte, microglia, BMEC, and pericyte after OGD/R. Six PH-associated microRNAs (miRNA-126a-5p, miRNA-132-3p, miRNA-142-3p, miRNA-195-3p, miRNA-29c-3p, miRNA-218a-5p) had significantly differentially expression in the model of BMEC or pericyte after OGD/R.

Recent studies suggested that the pathophysiology of cellular excitotoxicity, cell death signaling, oxidative stress and mitochondrial dysfunction, neuroinflammation, and BBB disruption play a pivotal role in cerebral ischemia. Immune cells in the brain and peripheral blood participate in acute and subacute pathogenesis of ischemic stroke. Therapies targeting signaling pathways involved in these molecular mechanisms provide the potential strategy for treating ischemic stroke and alleviating ischemic brain injury [42–44]. Modulating inflammatory and immune responses may be a potential crucial intervention to alleviate BBB disruption and HT in ischemic stroke [45]. Our study confirmed that 31 predicted hub target genes related to oxidative stress, apoptosis, immune, and inflammation may be associated with PH from the MCAO model after mechanical reperfusion. Twenty genes were involved in the mechanism of immune and or inflammation. Twelve genes were associated with all the oxidative stress, apoptosis, immune, and inflammation. Twelve genes had been reported as a potential regulator for HT in ischemic stroke in previous studies, including CASP3, CSF2, CTNNB1, CXCL12, HIF1A, IL1B, MTOR, PTGS2, PTPRC, TLR2, TLR4, VCAM1 [38, 39, 46–55]. The other 19 genes may be a potential therapy target for improving PH in acute ischemic stroke after mechanical reperfusion treatment.

Valuable microRNAs can regulate several target genes involved in multiple PH-related mechanisms. Simultaneously differentially expressed PH-associated microRNAs in neurons, astrocytes, microglia, BMEC, and pericytes under ischemic-reperfusion conditions may serve as therapeutic targets to alleviate ischemic brain injury. Recent studies showed that miRNA-218-5p expression was increased in astrocytes during OGD/R. Inhibition of miRNA-218-5p reduced inflammation, oxidative stress, and apoptosis damage induced by ischemic reperfusion injury [56, 57]. In our study, the differential expression of miRNA-218a-5p and its two key predicted target genes of IL1B and TLR4 were found in the peri-infarct of rat PH model and in-vitro OGD/R model of neuron, astrocyte, microglia, and pericyte. Both IL1B and TLR4 were involved in all the immune, inflammation, oxidative stress, and apoptosis mechanisms. MiRNA-195-5p was significantly downregulated during cerebral ischemic reperfusion injury in the OGD/R model of BMEC and the MCAO model. Increased expression of miRNA-195-5p alleviated cerebral ischemic reperfusion injury [58]. In our study, miRNA-195-3p was up-regulated, and its three key predicted target genes of CREB1, FOXO1, and STAT1 were decreased in the peri-infarct of the rat PH model. The differential expression of miRNA-195-3p and these predicted target genes were found in the BMEC and pericyte after OGD/R. CREB1 is involved in the mechanism of intracerebral hemorrhage by regulating inflammation and BBB disruption [59]. FOXO1 may be involved in the OGD/R-induced injury in the BMEC and play a role in the early brain injury after subarachnoid hemorrhage [60, 61].

This study has some limitations. We assessed the expression of HT-related microRNAs in the neuron, astrocyte, microglia, BMEC, and pericyte after OGD/R, respectively. However, each microRNA may be involved in the cross-talk process among the interaction of these cells during cerebral ischemia-reperfusion. In addition, this study did not evaluate the direct interactions of these PH-related microRNAs with their target genes by dual luciferase reporter technology. Finally, the exact interventing effect of these differentially expressed microRNAs on the PH by regulating their target genes requires further study through the animal and OGD/R models.

## Conclusions

Fourteen microRNAs were associated with PH after mechanical reperfusion in the rat MCAO and in-vitro OGD/R model. Simultaneously differentially expressed microRNAs and the associated genes in the neuron, astrocyte, microglia, BMEC, and pericyte may serve as valuable targets for PH after endovascular mechanical reperfusion in AIS. The vital role of reperfusion-inducible microRNAs and their related target genes via the signaling pathway in PH after mechanical reperfusion requires further investigation.

## Funding statement

This study was funded by the National Natural Science Foundation of China (81720108014, 81371275) and the Science and Technology Planning Project of Guangzhou City (201704020166).

## Data availability statement

The datasets of supplementary material for this article can be acquired from the corresponding author.

## Ethics statements

The study was reviewed and approved by the Institutional Animal Care and Use Committee of Sun Yat-sen Memorial Hospital, Sun Yat-sen University.

## Author contribution statement

Z-SS participated in the design of the present study. All authors participated in the interpretation and collection of the data. J-KZ, Z-RH, WQ and Z-SS wrote the initial manuscript. Z-SS revised the manuscript. All authors critically reviewed and edited the manuscript and approved the final version.

## Conflict of interest statement

The authors declare no conflict of interest.

---

**Supplementary Table S1.** Sequences of the primer for mature microRNAs used in the qRT-PCR.

**Supplementary Table S2.** Differential expression of parenchymal hematoma-related microRNAs in both peri-infarct and infarction core in the rat model.

**Supplementary Table S3.** Differential expression of parenchymal hematoma-related microRNAs in neuron, astrocyte and brain microvascular endothelial cell model with oxygen-glucose deprivation/reoxygenation (OGD/R).

**Supplementary Table S4.** Target genes for 14 PH-associated microRNAs predicted by TargetScan or miRDB databases.

**Supplementary Table S5.** The top ten enriched terms of molecular function, biological process, cellular component of GO, and KEGG pathways.

**Supplementary Table S6.** Predicted target genes involved in the mechanism of immune, inflammation, oxidative stress, and apoptosis.

**Supplementary Table S7.** The top twenty hub genes related to immune, inflammation, oxidative stress, and apoptosis were calculated by the MCC method in the CytoHubba plugin.

**Supplementary Table S8.** Forty-six key predicted target genes related to immune, inflammation, oxidative stress, and apoptosis for 14 PH-related microRNAs.

**Supplementary Table S9.** Differential expression of 46 predicted hub genes associated with parenchymal hematoma.

**Supplementary Table S10.** The microRNA-mRNA regulatory axes of parenchymal hematoma based on the prediction tool and PCR examination.

**Supplementary Figure S1.** Dynamic expression of 14 microRNAs in neurons after oxygen-glucose deprivation/reoxygenation (OGD/R) at 0, 6, 12, 24, 48, 72 hours.

**Supplementary Figure S2.** Dynamic expression of 14 microRNAs in astrocytes after oxygen-glucose deprivation/reoxygenation (OGD/R) at 0, 3, 6, 12, 24 hours.

**Supplementary Figure S3.** Dynamic expression of 14 microRNAs in microglia after oxygen-glucose deprivation/reoxygenation (OGD/R) at 0, 3, 6, 12, 24 hours.

**Supplementary Figure S4.** Dynamic expression of 14 microRNAs in brain microvascular endothelial cells after oxygen-glucose deprivation/reoxygenation (OGD/R) at 0, 3, 6, 12, 24 hours.

**Supplementary Figure S5.** Dynamic expression of 14 microRNAs in pericytes after oxygen-glucose deprivation/reoxygenation (OGD/R) at 0, 3, 6, 12, 24 hours.

## Abbreviations

ACTB: actin, beta
AIS: acute ischemic stroke
AGT: angiotensinogen
BBB: blood-brain barrier
BCL2: B-cell lymphoma 2
BCL2L1: BCL2-like 1
BMEC: brain microvascular endothelial cell
CASP3: caspase 3
CCL5: chemokine C-C motif ligand 5
CCND1: cyclin D1
CD40: CD40 molecule
CD44: CD44 molecule
CD80: CD80 molecule
CREB1: cAMP responsive element binding protein 1
CSF1: colony stimulating factor 1
CSF1R: colony stimulating factor 1 receptor
CSF2: colony stimulating factor 2
CTNNB1: catenin beta 1
CXCL1: C-X-C motif chemokine ligand 1
CXCL10: C-X-C motif chemokine ligand 10
CXCL12: C-X-C motif chemokine ligand 12
CXCR4: C-X-C motif chemokine receptor 4
DMEM: Dulbecco’s modified Eagle medium
ECM: endothelial cell medium
ESR1: estrogen receptor 1
EVT: Endovascular thrombectomy
FOS: Fos Proto-Oncogene, AP-1 Transcription Factor Subunit
FOXO1: forkhead box O1
FOXP3: forkhead box P3
GO: Gene Ontology
HIF1A: hypoxia inducible factor 1, alpha subunit
HT: hemorrhagic transformation
IDO1: indoleamine 2,3-dioxygenase 1
IFNB1: interferon beta 1
IGF1: insulin-like growth factor 1
IL13: interleukin 13
IL15: interleukin 15
IL17A: interleukin 17A
IL1A: interleukin 1 alpha
IL1B: interleukin 1 beta
IL2: interleukin 2
JUN: jun proto-oncogene
KEGG: Kyoto Encyclopedia of Genes and Genomes
LEP: leptin
MCAO: middle cerebral artery occlusion
MTOR: mechanistic target of rapamycin kinase
OGD/R: oxygen-glucose deprivation/reoxygenation
PH: parenchymal hematoma
PPARG: peroxisome proliferator-activated receptor gamma
PTEN: phosphatase and tensin homolog
PTGS2: prostaglandin-endoperoxide synthase 2
PTPRC: protein tyrosine phosphatase, receptor type, C
SIRT1: sirtuin 1
STAT1: signal transducer and activator of transcription 1
TLR2: toll-like receptor 2
TLR4: toll-like receptor 4
TP53: tumor protein p53
VCAM1: vascular cell adhesion molecule 1

## Notes

### Competing Interest Statement

The authors have declared no competing interest.

